# A comprehensive study of metabolite genetics reveals strong pleiotropy and heterogeneity across time and context

**DOI:** 10.1101/461848

**Authors:** Apolline Gallois, Joel Mefford, Arthur Ko, Amaury Vaysse, Markku Laakso, Noah Zaitlen, Päivi Pajukanta, Hugues Aschard

## Abstract

Genetic studies of metabolites have identified thousands of variants many of which are associated with downstream metabolic and obesogenic disorders. However, these studies have relied on univariate analyses, reducing power and limiting context specific understanding. Here we aim to provide an integrated perspective of the genetic basis of metabolites by leveraging the Finnish Metabolic Syndrome In Men (METSIM) cohort, a unique genetic resource which contains metabolic measurements across distinct timepoints as well as detailed information on statin usage. We increase effective sample size by an average of two-fold by applying the Covariates for Multi-phenotype Studies (CMS) approach, identifying 588 significant SNP-metabolite associations, including 248 novel associations. We further show that many of these SNPs are master metabolic regulators, balancing the relative proportion of dozens of metabolite levels. We then identify the first associations to changes in metabolic levels across time as well as evidence of genetic interaction with statin use. Finally, we show an overall decrease in genetic control of metabolic processes with age.

---

The human metabolome includes over 100,000 small molecules, ranging from peptides and lipids, to drugs and pollutants^1^. Because metabolites affect or are affected by a diverse set of biological processes, lifestyle and environmental exposures, and disease states,^2^ they are routinely used a biomarkers^3^. Thanks to recent technological advances, diverse components of the metabolome are being measured in large human cohorts, offering new opportunities to improve our understanding of the molecular mechanisms underlying metabolism and corresponding human traits and diseases^4^. For example, previous work has highlighted the role of metabolites in diseases such as Type 2 Diabetes^5,6^, cardiovascular disease^7^, and obesity^3,8^. Here we focus on the identification of genetic variants with pervasive effects on the metabolome, and those with effects dependent on statin treatment and age, two established modifiers of metabolite profiles^9^ and disease risk. Our study also introduces several analytical novelties. First, unlike previous genetic analyses of metabolites^10–17^, we leverage the high correlation structure between metabolites to increase the power via the CMS method^18^. Second, we used analytical and graphical tools to produce an integrated view of the genetic-metabolite network. Third, we use bivariate heritability and interaction analyses to examine changes in genetic regulation of metabolites as a function of aging and exposure to statins.

We first performed genome-wide association studies (GWAS) of 158 serum metabolites measured with nuclear magnetic resonance (NMR) in 6,263 unrelated individuals from the METSIM^19^ cohort. These measurements consisted of 98 lipoproteins (42 VLDL, 7 IDL, 21 LDL and 28 HDL), 9 amino acids, 16 fatty acids, and 35 other molecules (**Supplementary Table 1** and **Supplementary Figure 1**). GWAS was performed using standard linear regression (*STD*), but also using the *CMS* approach^18^, a powerful method we recently developed for the analysis of multivariate datasets (**Online Methods**). For both methods we tested association between each SNP and each metabolite while adjusting for potential confounding factors, including age and medical treatments (statins, beta blockers, diuretics and fibrate). We grouped significant SNP within independent linkage disequilibrium blocks and obtained 588 locus-metabolite associations involving a total of 54 loci (**Supplementary Table 3-4**). **Figure 1a** shows that these associations are spread over the 158 metabolites: we found 399 associations with lipoproteins (189 with VLDL, 38 with IDL, 88 with LDL and 84 with HDL), 17 with amino acids, 50 with fatty acids, and 122 with other molecules. Among these associations, 9 were significant with STD only (1.53%), 261 with both STD and CMS (44.39%) and 318 (54.08%) with *CMS* only (**Supplementary Table 5**). Overall, CMS led to a 118% increase in identified signals (**Supplementary Figures 2-3**). Among the 588 locus-metabolite associations identified, 248 signals (involving 34 loci) were not identified at the genome-wide significant level by previous large-scale metabolite studies. As illustrated in **Figure 1b**, new associations exist for 110 of the 158 metabolites. Among the 248 signals, 3 were significant with *STD* only (1.21%), 176 with both STD and *CMS* (70.97%) and 69 (27.82%) with *CMS* only. For each new association, we further mapped the top SNPs per locus to their nearest gene in a window of 100kb. **Table 1** presents the aggregated results, while complete details are provided in **Supplementary Table 6**.

**Figure 1:**
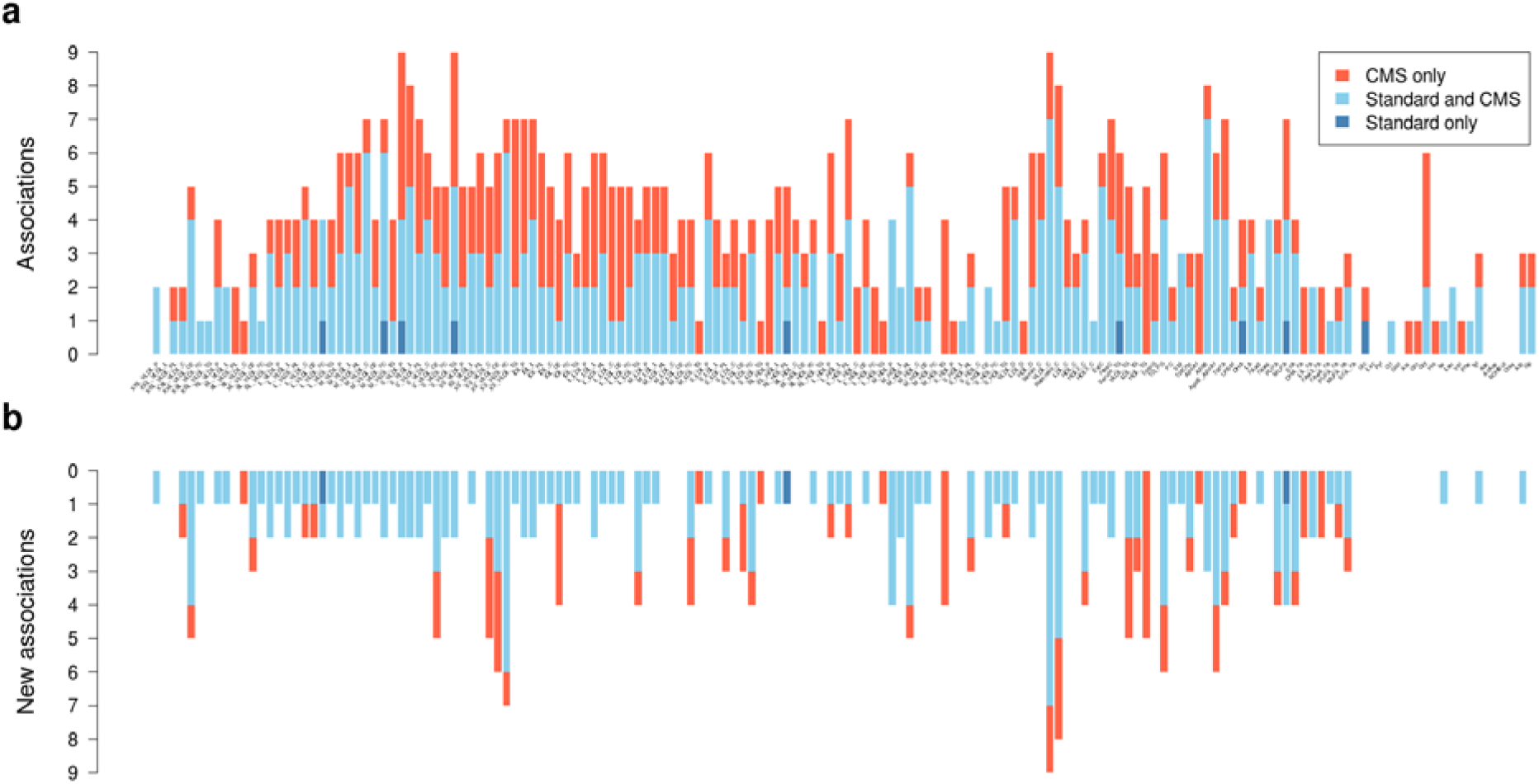
Locus - metabolite associations distribution. Distribution of the 588 significant associations (P < 1.28 x 10^−9^) identified in the 158 metabolites GWAS in the METSIM cohort. (a) Loci in dark blue were significant for standard linear regression adjusted by confounding factors. Loci in red were significant for linear regression adjusted with confounding factors and covariates selected by *CMS*. Loci in light blue were significant for both models. (**b**) Same plot including only the 248 new associations, not identified in previous metabolites GWAS (P > 1.28 x 10^−9^ in Teslovich^16^, Kettunen^15^, Rhee^17^ and Shin^13^ studies)

**Table 1:** New gene – metabolite associations.

| Chr. | Gene<sup>1</sup> | Position | SNP<sup>2</sup> | A1 | A2 | Associated metabolites | Opposite association |
| --- | --- | --- | --- | --- | --- | --- | --- |
| 1 | CELSR2 | 109818530 | rs646776 | T | C | ApoB_ApoA1, M_LDL_FC, S_LDL_FC/PL |  |
| 1 | DOCK7 | 63056112 | rs1748197 | G | A | HDL_TG, MUFA, M_HDL_P/PL/TG, PC, PUFA, TotCho, TotFA, XXL_VLDL_CE |  |
| 1 | GALNT2 | 230294916 | rs2144300 | C | T | ApoB, L_VLDL_*, M_VLDL_*, S_VLDL_FC/L/P/PL/TG, TG_PG, VLDL_C/D/TG, XL_VLDL_P/TG | M_HDL_PL, S_HDL_PL |
| 1 | PCSK9 | 55505647 | rs11591147 | G | T | ApoB_ApoA1, IDL_CE, L_LDL_TG, M_LDL_FC, Remnant_C, S_LDL_FC, S_VLDL_CE, VLDL_C, XL_HDL_FC, XS_VLDL_C/CE/FC, XXL_VLDL_CE | M_HDL_C/CE/P/PL, S_HDL_PL |
| 1 | PSRC1 | 109822166 | rs599839 | A | G | S_LDL_CE |  |
| 2 | APOB | 21225281 | rs1042034 | T | C | TotFA |  |
| 2 | GCKR | 27730940 | rs1260326 | T | C | Remnant_C, TG_PG, VLDL_C/TG, XL_VLDL_CE/FC | L_HDL_PL |
| 3 | PROK2 | 71880578 | rs7622817 | G | A | Serum_C |  |
| 4 | UTP3 | 71552398 | rs16845383 | A | G | Alb |  |
| 5 | MARCH3 | 126267351 | rs12655258 | C | T | HDL2_C |  |
| 5 | MIR4634 | 174223234 | rs12660057 | G | A | M_HDL_L |  |
| 6 | MICB | 31236410 | rs34131062 | T | C | S_VLDL_TG, VLDL_TG, XS_VLDL_TG |  |
| 6 | MIR3925 | 36613812 | rs6457931 | G | T | XL_HDL_L |  |
| 8 | LPL | 19832646 | rs17482753 | G | T | ApoB, HDL_TG, MUFA, SFA, TG_PG, TotFA, VLDL_C/TG |  |
| 8 | TRIB1 | 126485531 | rs7846466 | T | C | L_VLDL_C/CE/FC/L, MUFA, Remnant_C, VLDL_C, XL_VLDL_C/CE/L, XXL_VLDL_C/CE/FC |  |
| 10 | PCDH15 | 56015656 | rs11004183 | G | A | IDL_C/FC/L/P |  |
| 10 | PKD2L1 | 102075479 | rs603424 | G | A | MUFA_FA |  |
| 11 | APOA5 | 116660686 | rs2266788 | G | A | ApoB_ApoA1, HDL_TG, lle, M_HDL_TG, PUFA, Remnant_C, SFA, S_VLDL_CE, TG_PG, VLDL_C/TG, XS_VLDL_FC, XXL_VLDL_C/CE | HDL_D, L_HDL_P |
| 11 | CELF1 | 47539697 | rs4752845 | T | C | ApoA1, L_HDL_PL | XXL_VLDL_P |
| 11 | FADS1 | 61569830 | rs174546 | C | T | EstC, Faw3_FA, UnSat, XS_VLDL_L | M_VLDL_FC |
| 11 | FADS2 | 61597972 | rs1535 | G | A | DHA_FA, SM, XS_VLDL_FC | LA_FA, M_VLDL_P, XL_VLDL_TG |
| 11 | FADS3 | 61639573 | rs174448 | G | A | M_VLDL_PL |  |
| 11 | MTCH2 | 47663049 | rs10838738 | G | A | TG_PG |  |
| 11 | MYRF | 61551356 | rs174535 | C | T | PUFA_FA | S_HDL_TG |
| 11 | PTPMT1 | 47583121 | rs12798346 | C | T | HDL_D, L_HDL_P, XL_HDL_PL |  |
| 11 | TMEM258 | 61557803 | rs102275 | C | T | MUFA, MUFA_FA | HDL2_C |
| 12 | HNF1A | 121420260 | rs7979473 | G | A | M_LDL_P |  |
| 15 | LIPC | 58683366 | rs1532085 | A | G | HDL2_C, HDL3_C, HDL_TG, IDL_CE, LDL_TG, L_HDL_TG, L_LDL_L/TG, MUFA_FA, M_HDL_L/P/PL/TG, M_LDL_L/TG, PUFA, Remnant_C, SFA, S_HDL_TG, S_LDL_TG, S_VLDL_C/CE/FC/L/P/PL, TotCho, VLDL_C, XS_VLDL_C/CE/FC | Faw6_FA, LA_FA, PUFA_FA |
| 15 | LIPC-AS1 | 58730498 | rs588136 | C | T | LDL_TG, L_LDL_L, M_LDL_L/TG |  |
| 15 | LOC283665 | 58380442 | rs12910902 | T | C | IDL_TG, LDL_TG, L_HDL_CE/L, L_LDL_TG |  |
| 15 | MYO1E | 59453384 | rs2306791 | T | C | S_LDL_P/PL |  |
| 16 | C16orf47 | 73177225 | rs9673570 | A | G | Tyr |  |
| 16 | CETP | 56991363 | rs183130 | C | T | ApoB, ApoB_ApoA1, HDL_TG, IDL_L/P/PL, L_LDL_C/CE/L/PL, M_HDL_TG, Remnant_C, S_VLDL_CE, VLDL_C, XL_VLDL_CE, XS_VLDL_C/CE/FC, XXL_VLDL_CE | HDL2_C |
| 16 | DHX38 | 72144174 | rs9302635 | T | C | SFA, TotFA |  |
| 16 | ITGAM | 31343769 | rs4597342 | T | C | TG_PG |  |
| 16 | PMFBP1 | 72230112 | rs9923575 | T | C | UnSat |  |
| 19 | APOC1 | 45415640 | rs445925 | G | A | IDL_CE, M_LDL_FC, S_HDL_CE, S_LDL_CE/FC/PL, TotCho, XS_VLDL_C/CE |  |
| 19 | APOE | 45408836 | rs405509 | G | T | M_HDL_P/PL, PUFA |  |
| 19 | LDLR | 11202306 | rs6511720 | G | T | ApoB_ApoA1, IDL_CE, LDL_TG, L_LDL_TG, M_LDL_FC, Remnant_C, S_LDL_CE/FC, S_VLDL_CE, XS_VLDL_C/CE/FC |  |
| 19 | NECTIN2 | 45373565 | rs395908 | G | A | ApoB_ApoA1, Remnant_C |  |
| 19 | PRKCSH | 11560347 | rs755000 | T | G | FreeC |  |
| 19 | TOMM40 | 45395266 | rs157580 | A | G | VLDL_C, XS_VLDL_FC |  |
| 20 | PLTP | 44545048 | rs4810479 | C | T | S_HDL_FC/PL |  |

Chr., chromosome;
<sup>1</sup>Nearest gene from SNP, in a window of 100 kb before and after the SNP<sup>2</sup>SNP strongly associated with the majority of phenotypes present in last two columns, most significant SNP for each phenotypes are listed in **Supplementary Table 6**

We next performed in silico replication for all new association signals using data from four previous metabolites GWAS^13, 15–17^ (**Supplementary Table 2**). Note that the metabolites analyzed differ widely across the replication datasets, and there was only a partial overlap with the METSIM’s metabolites. In practice, the bulk of the replication analysis was performed using data from Kettunen et al^15^ (N=24,925), while the other studies were informative only for a very limited set of metabolites. Furthermore, we focused replication only in the subset of overlapping metabolites. Out of the 248 new SNP-metabolites pairs, 102 were available for in-silico replication (41.1%). Among those, 73 (71.6%) were replicated at a nominal threshold of 5%. Finally, when comparing the top SNPs from every loci associated with at least one metabolite (N=70, see next paragraph) with previous GWAS on coronary heart disease (CHD)^16^, body mass index (BMI)^17^ and type 2 diabetes (T2D)^18^, we observed substantial enrichment for nominally significant association. Given a false discovery rate (FDR) at 10%, we observed 30 significant genes for CHD, 5 for BMI and 4 for T2D (in bold in Supplementary Table 7), indicating that part of these variants are also likely involved in the genetics of these common diseases.

We observed substantial polygenicity and pleiotropy. Using the aforementioned SNP-gene assignment, 147 metabolites were associated with at least one gene, and a total of 70 genes associated with at least one metabolite. Metabolites were associated with 1 to 9 genes, with an average of 4 genes. On the other hand, genes showed high level of pleiotropy with an average of 8.4 metabolites associated with each gene. Although, 13 genes (LIPC, APOA5, CETP, PCSK9, LDLR, GCKR, APOC1, LPL, GALNT2, CELSR2, TRIB1, DOCK7 and FADS2) capture over 75% (N=457) of all associations (**Supplementary Figure 4**). These extensive pleiotropic effects are illustrated in **Figure 2**, which includes all associations plotted in a Cytoscape^20^ network. The network highlights several known master regulatory effects of genes. For example, *CETP* encodes a protein that transports cholesterol esters and triglycerides between HDL metabolites and VLDL metabolites. Our network clearly displays the opposite effect of variants in *CETP* on HDL and VLDL. Our results also contribute explaining the complex effect of PCSK9. Besides its established association with LDL and VLDL, our analyses confirm opposite associations with HDL metabolites^21^. Overall, the gene displaying the strongest pleiotropic effect was *LIPC* with 75 associated metabolites, of which 34 were new associations (11 of them were available for replication, and 8 were replicated at a 5% alpha threshold).

**Figure 2:**
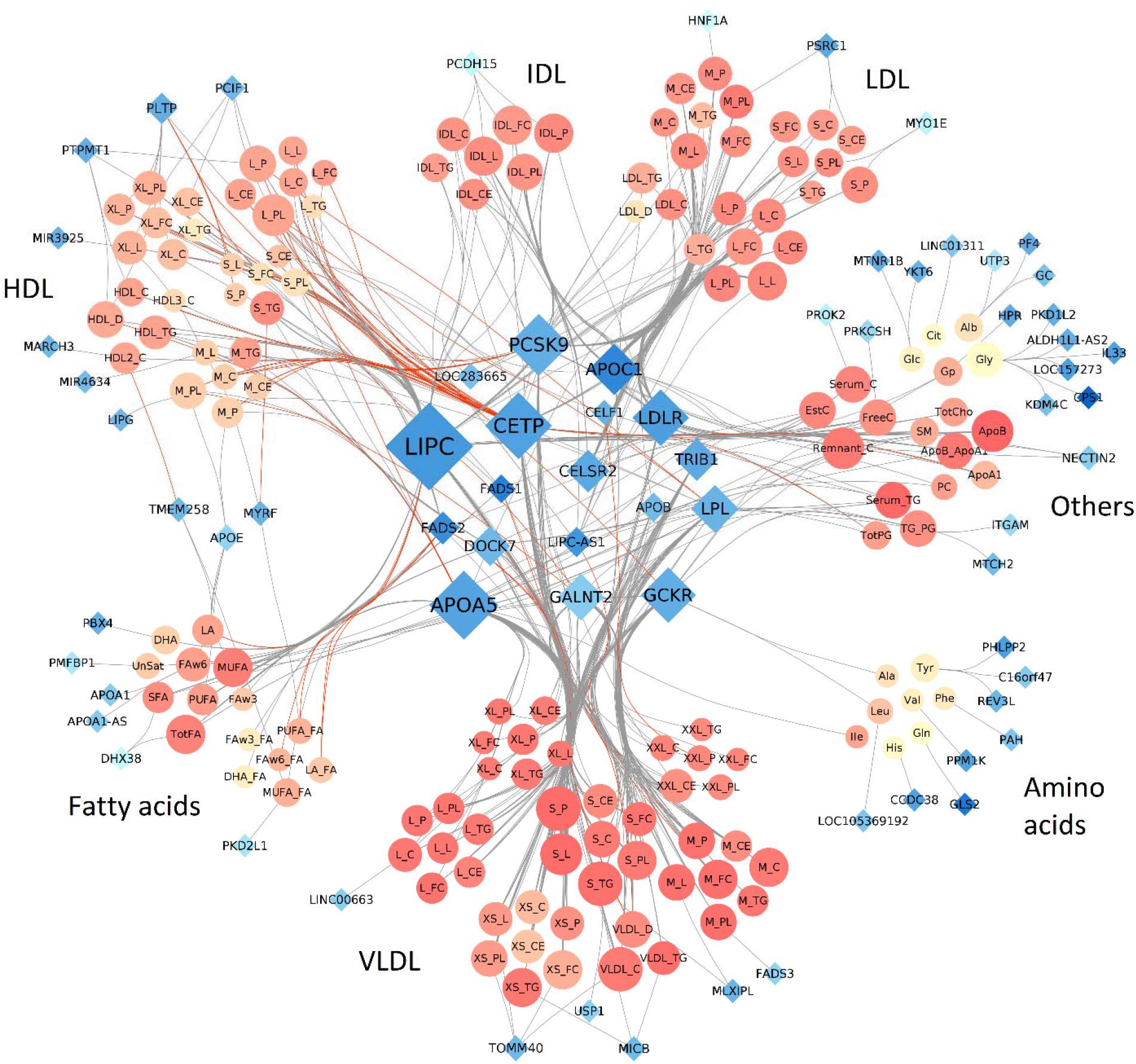
Network representation of locus-metabolite associations. Network representation of the 588 locus-metabolite associations identified in the 158 metabolites GWAS in METSIM. Each node represents either a gene (blue diamonds, 70) or a metabolite (orange circles, 147). Each edge is an association between one gene and one metabolite. Node size is directly proportional to the number of nodes associated with it. Red edges correspond to opposite effect of a gene on a metabolite, compared to the other metabolites associated with the same gene. Metabolites colors (orange shades) represents correlation strength between a given metabolite and all other metabolites. Genes colors (blue shades) represent strength of correlation between a given gene and associated metabolites, quantified as the average of *r*-squared across all corresponding metabolites.

To illustrate how multivariate results can help identify likely causal variants, we then applied the FINEMAP^22^ algorithm to the 75 metabolites associated with the latter *LIPC* region (**Supplementary Note** and **Supplementary Table 8**). Our analysis identified 3 distinct association signals (**Figure 3** and **Supplementary Tables 9-10**) with consistently high probabilities of causal effect on triglyceride in HDL and LDL from 7 SNPs. We cross-referenced top variants of these three signals with GWAS of common human diseases^23^, and functional annotations from *Haploreg*^24^. The first signal is composed only of SNP rs10468017, which was previously strongly associated with age-related macular degeneration^25,26^, but also with cardiovascular diseases^27^ and metabolic syndrome^28^. It lives in a region harbouring H3K4me1/H3K4me3 and H3K27ac/H3K9ac marks of promoter and enhancer in Adipose Derived Mesenchymal Stem Cell Cultured Cells. The high number of rs10468017-metabolite associations in our study, and previous analyses^29^ suggests an overall effect of rs10468017 on *LIPC* expression. The second signal includes 4 SNPs in complete linkage disequilibrium that were previously associated advanced age-related macular degeneration^30^. It colocalizes with histone marks of promoters and enhancers in liver. These SNPs are also in a region bound by 4 transcription factors: FOXA1 (rs1077834); FOXA1 and FOXA2 (rs1800588); and RXRA and USF1 (rs2070895). Among those transcription factors, USF1 has been associated with low-density lipoprotein cholesterol levels, triglycerides^31,32^, and combined hyperlipidemia^33,34^. Furthermore, USF1 has been implicated in the expression of hepatic lipase^35^, making rs2070895 the strongest candidate for potential functional effects through differential regulation of *LIPC*. Finally, the last signal included 2 SNPs, among which rs113298164 clearly harboured the highest number of relevant bio-features. It is a rare missense mutation in a region having promoter histone marks in hESC Derived CD184+ Endoderm Cultured Cells. The SNP is also detected by GERP^36^ as part of a sequences that is constrained across mammalian genomes. It induces a T405M mutation in LIPC protein and is referenced as involved in hepatic lipase deficiency^37^.

**Figure 3:**
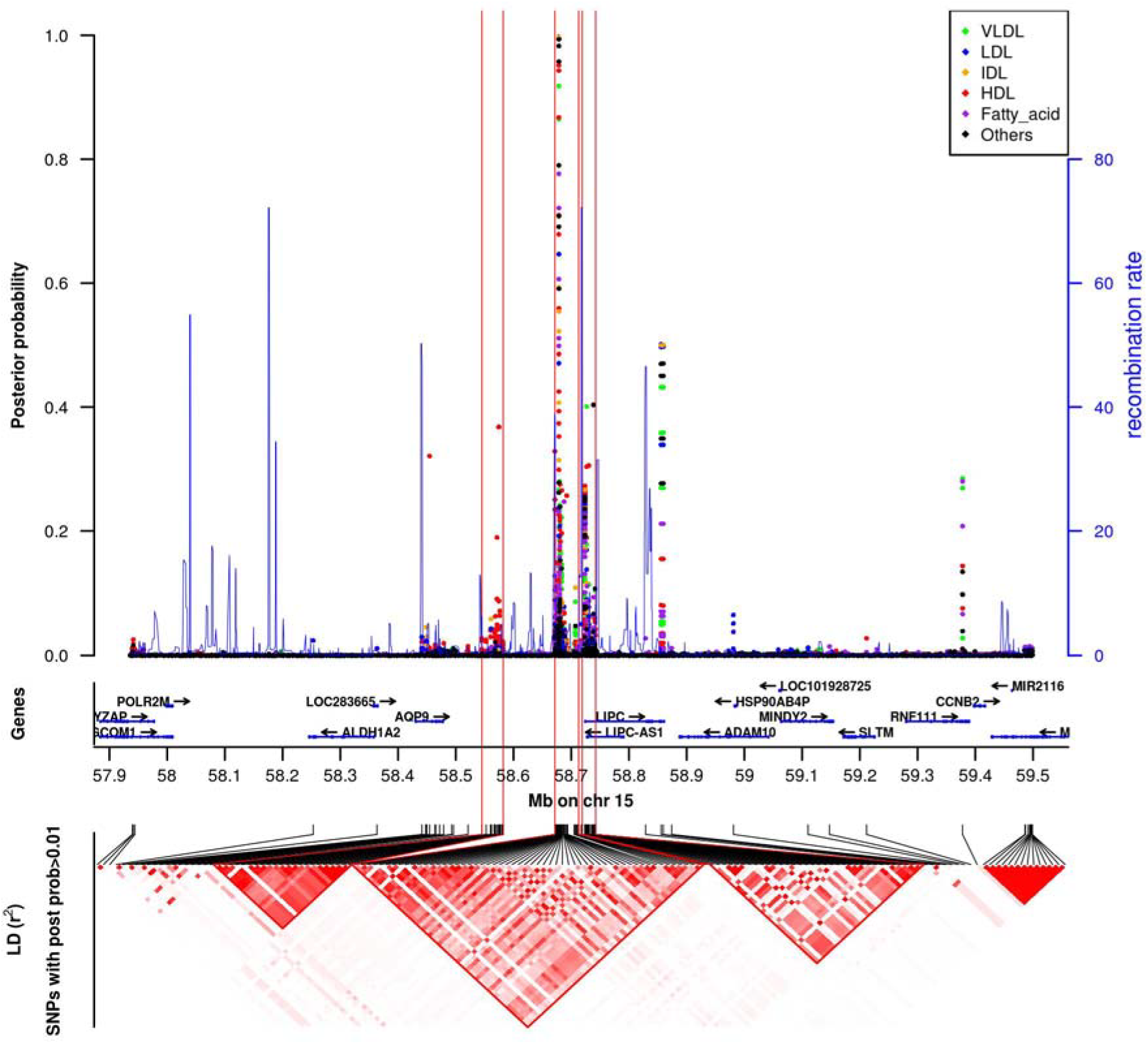
Regional plots of LIPC locus fine mapping. Panel (**a**) indicates the posterior probability assessing the evidence that the SNP is causal for each of the 75 phenotypes and the local recombination rate. Panel (**b**) contains genes from the UCSC hg19 annotation. Panel (**c**) is a r^2^-based LD heatmap computed using PLINK1.9 on the METSIM data. The gradient of red is proportional to the r^2^. For clarity, we represented the LD only for SNPs with a posterior probability >0.01 for at least 1 phenotype.

An important component of the METSIM cohort is the collection of statin use amongst participants. To examine changes in genetic regulation of metabolites when taking statins, we performed an interaction test between SNPs and statin for each of the 588 locus-metabolite associations. While no interaction test passed a Bonferroni correction for multiple testing (i.e., p < 8.5 x 10^−5^, **Supplementary Table 11**), 83 out of the 588 locus-metabolite association showed nominally significant interactions (*p*-value < 0.05). Based on the q-value distribution^38^, there were 35 significant interactions at a 10% FDR, showing that at least some of the identified SNP-metabolite effects depends on statin use status. Many of these 35 interactions involve the same two genes, TRIB1 (associated with VLDL particles) and APOC1 (mostly associated with LDL and IDL particles), while other genes (FADS1, FADS2, MARCH3, MIR3925, MIR4634, and ITGAM) show interaction with a single metabolite. Interestingly, APOC1, is associated with statin-mediated lipid response^39^, and previous work suggests that FADS1 and FADS2 might modulate response to simvastatin^40^. We also checked statin interaction in follow-up data (see ***Online methods***), and found limited interaction values, except for APOC1 region, in which 90% of interaction signals found in baseline data are replicated.

Another unique aspect of the METSIM cohort is a second measurement of the same metabolites, using the same technology, approximately five years after the baseline (***Online methods***) for 3,351 unrelated individuals. We used these data to screen for genetic variants associated with an intra-individual change in metabolites level across time. In practice, we applied the same strategy as for our primary analysis but using the difference between follow-up and baseline data divided by age difference as outcome (Δ_*fb*_ = (*f* – *b*)/(*age_f_* – *age_b_*)), while adjusting for the same confounding factors as baseline and covariates selected by *CMS* in baseline measurements. There were 30 SNP-metabolites pairs reaching the standard 5 x10^−8^ *p*-value threshold with either *STD* or *CMS* (**Supplementary Table 12**), corresponding to 8 locus-metabolite associations (**Supplementary Table 13**). To the best of our knowledge, these are the first reported SNPs associated with changes in metabolic activity during aging. These associations involved 7 metabolites: S-HDL-TG, VLDL-C, DHA, DHA/FA, LA/FA, Faw3/FA, FAw6/FA, and 6 genes: *PDZRN4, LGMN, FADS1, FADS2, TNIK, LIPC*. Four of these associations were genome-wide significant in the marginal association at baseline (P < 5 x 10^−8^). The four new signals were observed for S_HDL_TG, VLDL_C, LA_FA and Faw6_FA. We also performed age interaction test on the linear regression between ∆_*bf*_ and significant SNPs (**Online methods**). However, none of the age interaction *p*-values was significant.

To examine global changes of genetic regulation of metabolites across time we also estimated heritability for each phenotype at each timepoint as well as the genetic and environmental correlations of the same phenotype between timepoints using bivariate linear mixed models^41,42^. **Figure 4** and **Supplementary Table 14** give heritability values for each metabolite, in both baseline and follow-up data. To avoid any bias in heritability estimation, we computed it on samples present in both baseline and follow-up studies and excluded those who were present in baseline study only. The average heritability decreased from 24.9% at baseline to 18.8% at follow up, with only 30.8% (p-value < 2e-9) having higher heritability at follow-up. The sample size was not large enough to estimate genetic correlation with low standard error, but the average estimate of 0.92, and the strong correlation of fixed effect sizes between time points (**Supplementary Table 15**), suggests that increasing environmental variance as opposed to decreased genetic variance underlie the reduction in heritability. If true, this result might also explain the absence of SNP-by-age interaction signal in our previous analysis.

**Figure 4:**
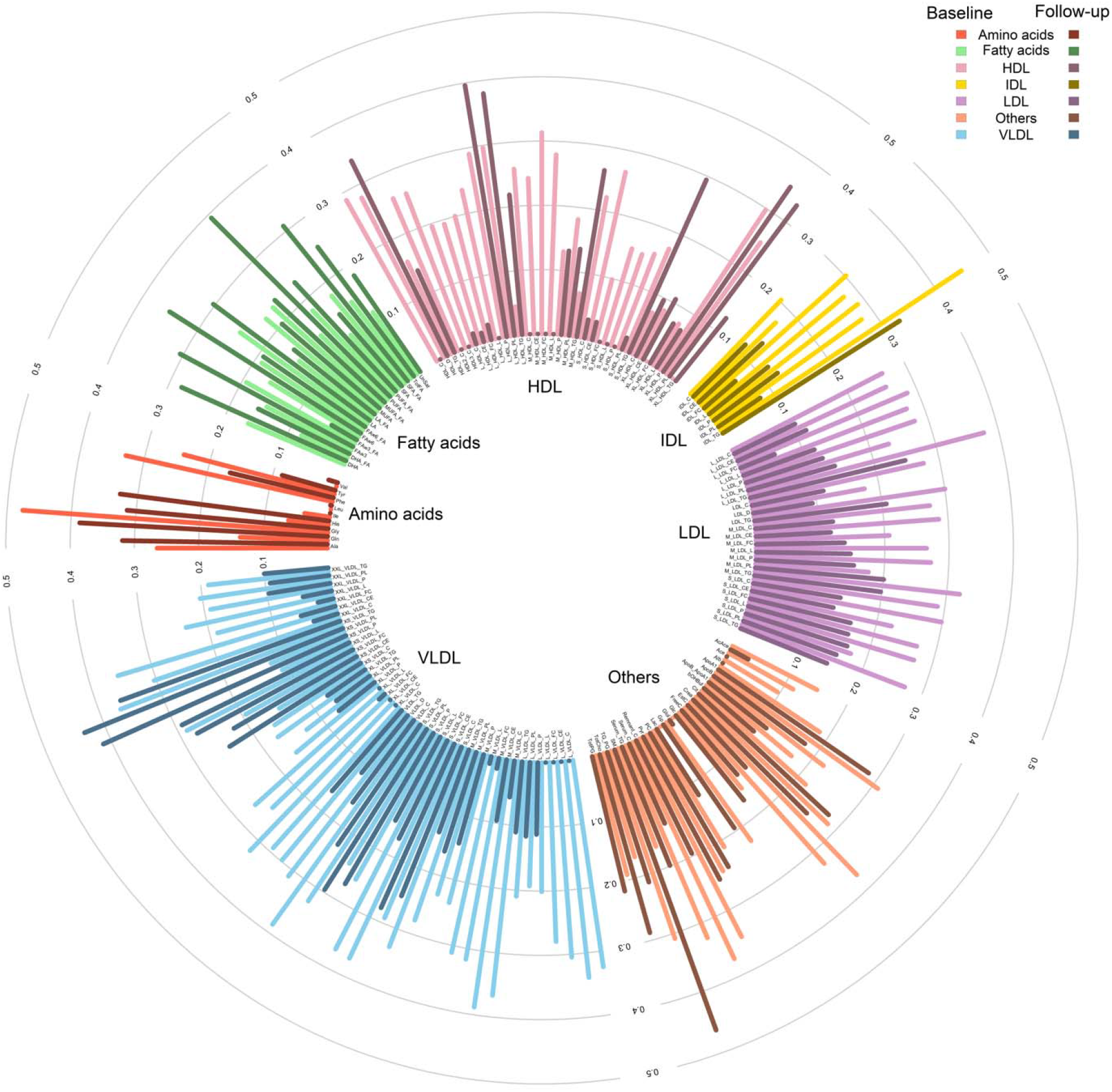
Heritability of metabolites in baseline and follow-up data. Heritability of studied metabolites, computed on individuals present in both baseline and follow-up data. We used bivariate restricted maximum likelihood (REML) and included 10 genetic PCs, age, and age as fixed effects. Light colors stand for heritability in baseline data and dark colors stand for follow-up data.

There are several shortcomings of this work. The study can be improved by adding the related individuals in the model, further increasing power. However, CMS cannot currently handle related individuals in reasonable computational time. The study can also be extended to imputed SNPs to improve fine-mapping estimates. TWAS estimates^43^ were not available for many of the core metabolic genes, but they could become feasible as larger RNA-seq data sets across more tissues are produced. Finally, direct perturbations of individual genes in cell lines or model organisms could help resolve the causal genes in the associated loci.

## Acknowledgements

We thank the METSIM individuals who participated in this study. This study was funded by National Institutes of Health (NIH) grants HL-095056, HL-28481, and U01 DK105561.

## URL

The code to run CMS on a large data set is available at https://gitlab.pasteur.fr/statistical-genetics/runCMS.

## ONLINE METHODS

### METSIM cohort

The METSIM cohort^19^ is composed of 10,197 Finnish men from 45 to 73 years old and aimed at investigating non-genetic and genetic factors associated with Type 2 Diabetes and cardiovascular diseases. Participants were recruited and examined between 2005 and 2010 in Kuopio town in Eastern Finland. The study was approved by the ethics committee of the University of Kuopio and Kuopio University Hospital, in accordance with the Helsinki Declaration. For each sample, 228 serum metabolites (lipids, lipoproteins, amino acids, fatty acids and other low molecular weight metabolites) measurements were made with nuclear magnetic resonance (NMR) at baseline. A follow-up study was conducted about 5 years after the baseline study. 6,496 participants (64 %) were reexamined with the same protocol and metabolites were measured a second time using the same technology. In our study, we considered 158 variables, including 150 raw measurements and 8 ratios. Other available variables, which were mostly percentages, were not included in the study. Besides metabolic measurements, several variables were also available including drug treatment and large group of other phenotypes. All samples were genotyped for 665,478 SNPs using the *Illumina OmniExpress* chip. Genotype data went through standard quality control, filtering individuals with missing rate below 5%, and SNPs with missing rate below 5% or with P < 10^−5^ in Hardy-Weinberg test.

### Data pre-processing

In order to remove outliers without reducing sample size, we first applied inverse normal rank-transformation on every analyzed metabolite. This was done using the *rntransform* function in R package GenABEL^44^. Because of potential confounding effect of statins use on metabolites, we excluded all statins users (1,722 individuals) when analyzing LDL, IDL, Apolipoprotein B and cholesterol. We also excluded fibrates users (25 individuals) when analyzing VLDL, IDL, triglycerides and chylomicron for similar reason. Finally, we removed all individuals with a genetic relationship coefficient larger than 0.05 and used only unrelated individuals. After filtering, there remained 6,263 samples available for analysis. For SNP data, we filtered variants with a minor allele frequency (MAF) lower than 1%. 609,262 SNPs remained after filtering.

### Genome-wide association screening

We used two different models in the analysis. First, we performed a standard linear regression (STD) between each metabolite (*Y*) and each SNP (*G*), adjusted for established confounding factors (***C***): age and medical treatments (statins, diuretics, fibrate and beta blockers):

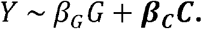

Then, we used the *CMS* algorithm to select additional covariates for each SNP-metabolite pair tested. Consider a metabolite *Y_k_*, which we refer further as the primary outcome. The *CMS* approach select potential covariates from the set of available metabolites *Y_l≠fk_*. In brief, the algorithm is divided in four steps. The first step is the computation of marginal effects through standard linear regressions between variables taken two by two: i) *Y_k_* ~ *G* where *G* is the genetic variant tested, ii) *Y_l≠k_* ~ *G* where *l* includes a subset of candidate covariates (see next paragraph) and iii) *Y_k_* ~ *Y_l≠k_*. The second step consists in filtering covariates based on a multivariate test between *G* and all *Y_l≠k_*. In practice, it uses a Multivariate analysis of variance (MANOVA), which is applied iteratively, removing one by one covariates potentially associated to the genetic variant tested, until *G* does not display association with Y_*l≠k*_ in the MANOVA. The third step is the filtering of covariates based on Y_*l≠k*_ ~ *G* association conditional on *Y_k_* ~ *G* association (see ***Supplementary Note***). It is a stepwise procedure that removes progressively covariates that are potentially associated with *G*. The last step consists in a linear regression between predictor and outcome, adjusted for the selected covariates (***Y***_*L*_):

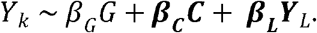

To address some of the limitations of *CMS*, we also applied for each outcome *Y_k_* a pre-filtering of candidate covariates *Y_l≠k_* before applying *CMS*. First, to avoid bias due to very high correlation between covariates and the outcome, we excluded all *Y_l≠k_* explaining more than 70% of the outcome variance. Second, to reduce the risk of false positive due to the inclusion of covariates that are hierarchical parent of the outcome under study, we excluded from the set of initial covariates all secondary outcome that were in the same biological group (LDL, HDL, …) as the primary outcome. Third, to reduce the computational burden, we reduced the number of candidate metabolites used as input of *CMS* to 30 through on AIC (Akaike information criteria, **Supplementary note, Supplementary Figures 5-6**). As showed in **Supplementary Figure 7**, it allows reducing substantially the computation time, while focusing on candidate covariates that altogether still explain a substantial proportion of the primary outcome variance.

### Post-GWAS processing

The threshold used to determine significant loci was calculated by dividing the standard genome wide significant threshold of 5 x 10^−8^ by the number of effective tests accounting for all variants tested and all metabolites. To estimate the number of effective tests, we first did a principal component analysis of our 158 metabolites. Then, we calculated the number of principal components that explained 99% of the total variance. We obtained 39 effective tests. The significance threshold was then 1.28×10^−9^.

Because of the great number of signals, we chose to summarize our results by loci, corresponding to approximately independent LD blocks. In practice, we sliced the genome in 1703 independent regions based on a recombination map recently described by Berisa et al^45^. These regions are 10 kb to 26 Mb long, with an average size of 1.6 Mb. For each region, we kept the SNP with the best p-value obtained by either *STD* or *CMS*. We then used the UCSC database to assign the closest gene to each SNP, with a maximum distance of 100 kb.

### GWAS of delta between baseline and follow-up across metabolites

We used data from baseline and follow-up studies to perform GWAS of the difference between the two time points, divided by the age difference. We called that variable ∆_*fb*_:

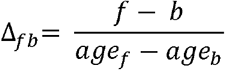

where *f* and *b* are metabolite measurements at follow-up and baseline, respectively. As for baseline data analysis, we used *STD* and *CMS* approaches, with covariates pre-selection based on AIC.

Confounding factors used for the baseline analysis were also included as covariate in all delta analysis. We did not adjust for baseline value in the main analysis.

### Interaction analyses

We performed two follow-up interaction analyses for subset of SNP-metabolite associations identified in the GWAS. First, we assessed SNP-by-age interaction effect in both baseline and follow-up analyses for the subset of SNP showing significant effects on delta in metabolite levels between baseline and follow-up (∆_*bf*_). In practice, we applied a standard linear regression between the corresponding outcome and genetic variant, adjusting for the same potential confounding factors as in the primary GWAS analysis, and adding the interaction term *β_int_G* ∗ *age*:

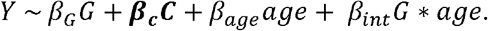

Second, we assess potential SNP-by-statin interaction for the 588 locus identified in the primary GWAS analysis. In that specific analysis, we included all statin users (which were removed in the primary analysis for some metabolites, as explained before) and performed linear regression between each metabolite and the best SNP in the associated loci (minimum p-value). The regression was adjusted by confounding factors and included the interaction term *β_int_ G* ∗ *statin*:

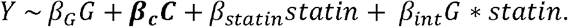

### Heritability

We first took a set of 3,342 individuals corresponding to the intersection between baseline and follow-up data. The baseline and follow up phenotypes were combined, normalized, and separated into baseline and follow up series, so the normalized phenotypes at baseline and follow up were directly comparable (i.e. equal normalized phenotypes at baseline and follow up correspond to equal raw phenotypes). We used GCTA’s bivariate REML^46^ and included 10 genetic PCs, age, and age^2^ as fixed effects. The effect sizes of the aformentioned fixed effects were strongly correlated at each time point (rho>0.6) and there were minimal differences in variance explained (<5%). Heritability estimates at the two time points were plotted using *circlize* R package^47^, while the complete GCTA output, including genetic and environmental variance estimates, genetic and environmental covariances, and LRT p-values for genetic correlation are provided in **Supplementary Table 14**.

